# One fly - one genome : Chromosome-scale genome assembly of a single outbred *Drosophila melanogaster*

**DOI:** 10.1101/866988

**Authors:** Matthew Adams, Jakob McBroome, Nicholas Maurer, Evan Pepper-Tunick, Nedda Saremi, Richard E. Green, Christopher Vollmers, Russell B. Corbett-Detig

**Affiliations:** Department of Molecular, Cellular, and Developmental Biology, University of California Santa Cruz, CA 95064; Department of Biomolecular Engineering, University of California Santa Cruz, Santa Cruz, CA 95064; UCSC Genomics Institute, University of California Santa Cruz, Santa Cruz, CA, 95064; Dovetail Genomics, Scotts Valley, CA, 95066

**Keywords:** genome, sequencing, assembly, Hi-C

## Abstract

A high quality genome assembly is a vital first step for the study of an organism. Recent advances in technology have made the creation of high quality chromosome scale assemblies feasible and low cost. However, the amount of input DNA needed for an assembly project can be a limiting factor for small organisms or precious samples. Here we demonstrate the feasibility of creating a chromosome scale assembly using a hybrid method for a low input sample, a single outbred *Drosophila melanogaster*. Our approach combines an Illumina shotgun library, Oxford nanopore long reads, and chromosome conformation capture for long range scaffolding. This single fly genome assembly has a N50 of 26 Mb, a length that encompasses entire chromosome arms, contains 95% of expected single copy orthologs, and a nearly complete assembly of this individual’s *Wolbachia* endosymbiont. The methods described here enable the accurate and complete assembly of genomes from small, field collected organisms as well as precious clinical samples.

## Introduction

The creation of high quality genome assemblies is a key step for the study of organisms on both the level of individuals and populations (Dudchenko et al. 2017). Conventional genome sequencing projects rely on whole-genome shotgun sequencing approaches that generate huge numbers of short sequence reads at low cost. While short reads can be reassembled into larger contiguous genome segments by identifying overlapping reads, they often fail to generate chromosome length assemblies due to the challenge of assembling repetitive DNA sequences. Consequently, many published genomes are highly fragmented (Worley et al. 2017). Fragmented genomes can be valuable for gene-level studies but many genomic analyses such as understanding chromosome-scale evolution, resolving full-length haplotypes, association studies, and quantitative trait locus mapping require high-quality chromosome-scale assemblies. New hybrid genome assembly approaches can produce highly contiguous assemblies that represent true chromosome length genomes (Rice and Green 2019).

Two recent advances in genomic technologies have dramatically raised the quality of genome assemblies (Yuan et al. 2017). First, third generation long-read sequencing technologies are capable of sequencing entire long repetitive sequences, but they suffer from higher error rates and lower throughput (Worley et al. 2017). Second, proximity-ligation sequencing, or Hi-C, produces short-read pairs representing sequences that are close together in three-dimensional space (Lieberman-Aiden et al. 2009). This allows high throughput “scaffolding” of challenging genomic regions (Putnam et al. 2016). However, these impressive gains in genome assembly quality have not been realized across all species due to important biological constraints.

Genome projects can be complicated by the small size of many organisms, which yield corresponding low amounts of DNA from a single individual. Consequently it is not always feasible to obtain sufficient input material for the genomic approaches described above without pooling individuals (Li et al. 2019). Nonetheless, developing applications for single individual genome assemblies offers several key advantages. First, it may not be possible to obtain more than a single individual for some species. Second, even if many could be found, pooling several individuals increases the genetic diversity in the DNA input, imposing challenges for accurate genome assembly. For wild caught samples, the possibility of combining cryptic species has the potential to impact assembly quality and introduce spurious biological conclusions. Finally, low input sequencing methods could be used to assemble genomes from precious clinical samples. There is therefore a clear need for new methods that can assemble highly contiguous genomes from a single isolate with limited available DNA.

Recently, Kingan et al. released a whole-genome assembly obtained from a single mosquito, *Anopheles coluzzii,* sequenced using three PacBio SMRT Cells (Kingan et al. 2019). Although the assembly has high contiguity (contig N50 3.5 Mb), the authors were unable to obtain chromosome-scale contigs or scaffolds and the resulting assembly does not include biologically important regions of the genome that contain chromosomal inversion breakpoints (Kingan et al. 2019; Corbett-Detig et al. 2019). Additionally, the input material used, approximately 100 ng of high quality DNA, may still be challenging to obtain from a single field-collected individual in many species. Nonetheless, this pioneering work suggests a powerful solution in developing low-input protocols for simultaneously obtaining Hi-C and long-read data from single individuals.

Here, we present a chromosome scale hybrid genome assembly of a single *Drosophila melanogaster* female. From this single individual, we produce long reads, short reads and proximity ligation sequencing data. Our assembly approach leverages the unique value added by each data type to produce a chromosome-scale and accurate genome assembly. This approach is applicable for millions of small species and for irreplaceable clinical samples.

## Results

### Sample Selection

Although numerous studies have assembled genomes from completely (Adams 2000) or partially (Kingan et al. 2019) inbred arthropods, the genomes of a field collected samples will likely be highly heterozygous outbred individuals. To make our assembly task conservatively challenging yet straightforward to evaluate, we generated an outbred fly by crossing females of the *D. melanogaster* reference strain *y; cn, bw; sp,* or ISO1 (Adams 2000), to males of another inbred and genetically distinct strain, I38 (Grenier et al. 2015). Importantly, I38’s genome is collinear with the reference on broad scales, although smaller rearrangements, such as small-scale indels and copy number variants, are almost certainly present in the genome (Grenier et al. 2015; Lack et al. 2015). We can therefore use progeny from this cross to demonstrate the applicability of our method for assembling genomes of outbred field-collected arthropod individuals and we can easily verify the accuracy of the assembly by comparison to the ISO1 reference genome. To facilitate the use of several sequencing methods, the single outbred fly chosen for sequencing (referred to as H3) was first laterally dissected (Figure 1).

**Figure 1.**
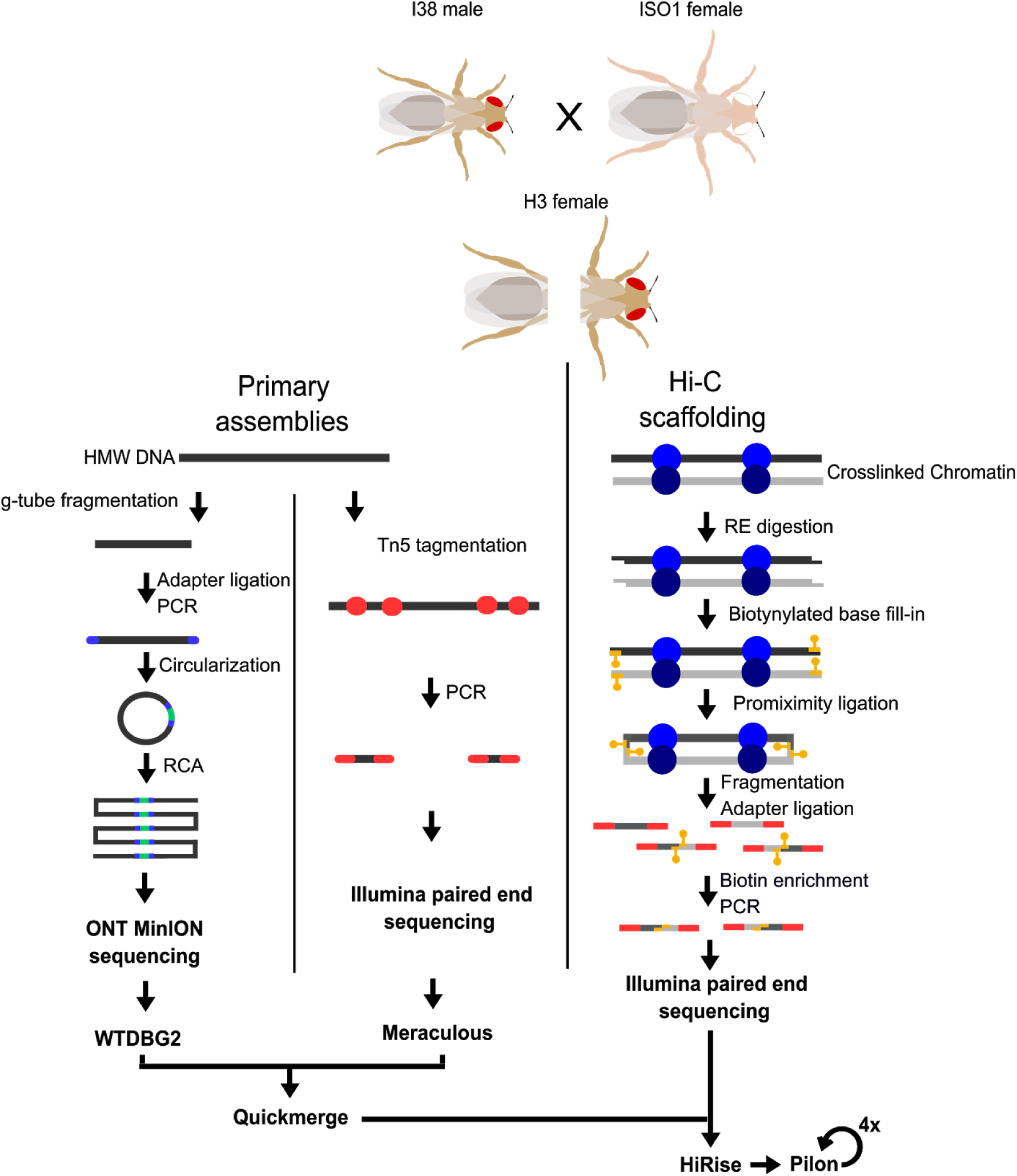
Experimental flow chart. A heterozygous fly (H3) was produced by crossing ISO1 and I38 strains. A single female offspring was laterally dissected. From the posterior half, HMW DNA was extracted and used to prepare the two primary assemblies, a R2C2 genomic library for nanopore sequencing, and a Tn5 tagmentation library for paired end Illumina sequencing. The anterior portion was used to isolate intact chromatin to generate a Hi-C paired end Illumina library. The two primary assemblies were merged into one then arranged into chromosome length scaffolds using the Hi-C contact frequency data.

### Primary Sequencing Datasets

From a single outbred adult female fly, we produced short-read shotgun, long-read shotgun and Hi-C libraries (Figure 1). From the posterior half, we extracted high molecular weight (HMW) DNA and we obtained approximately 104 ng in total. We used 78 ng to produce an Oxford Nanopore Technology (ONT) sequencing library following the R2C2 protocol (Volden et al. 2018) with slight modification for genomic DNA (see Methods). The R2C2 protocol generated ONT raw reads that contain tandem repeats of *Drosophila* genomic DNA sequence separated by splint sequences. The R2C2 post-processing pipeline (C3POa) processes these raw reads and generates two types of output reads: 1.) Consensus reads are generated if an ONT raw read is long enough to cover an insert sequence more than once which is evaluated by detecting a splint sequences in the raw read and 2.) Regular “1D” reads reads for which no splint could be detected in the raw read. In total, 277,305 consensus reads and 1,769,380 “1D” reads were generated from a single ONT MinION flow cell. Both read types were included in the assembly. We additionally produced an Illumina sequencing library using a standard Tn5-based protocol (Methods) and from this we obtained 133,135,777 total paired-end reads (Table 1).

**Table 1.** Summary of sequencing data used for assembly and scaffolding

| Library | Total Number of Reads | Read Length | Predicted Coverage |
| --- | --- | --- | --- |
| **Illumina Tn5** | 133,135,777 | 151 bp (paired end) | 333x |
| **ONT R2C2** | 2,046,685 | 3,541 bp (median length) | 60x |
| **Illumina HiC** | 68,400,787 | 151 bp (paired end) | 171x |

Because both R2C2 and our Tn5 protocol are optimized for low DNA inputs, they require some amplification to produce suitable quantities of libraries for high throughput sequencing. Likely as a consequence, the variance in sequencing depth exceeds the theoretically expected variance if reads were sampled uniformly at random from the genome. Indeed, for libraries with mean depths 236x and 39.7x we obtained depth variances of 8382 and 1038 for Tn5 and ONT respectively. Nonetheless, we show below that moderately long contigs can still be generated from these data (Supplementary Figure S1).

We also produced a Hi-C library to enable long-range scaffolding across the genome. We optimized a chromatin conformation capture sequencing method (Belton et al. 2012; Lieberman-Aiden et al. 2009) for application to samples with minimal input materials (See Methods). Using this approach and just the anterior half of the fly, we were able to produce 68,400,787 reads in total from a Hi-C library made from the thorax and abdomen (see Methods). This represents an average of approximately 93,991 clone coverage across the genome. Furthermore, despite low-input, the PCR duplication rate is quite modest (12%). These data therefore indicate that our single-fly Hi-C approach can produce high complexity libraries suitable for scaffolding high quality genomes.

### Primary assemblies

To accommodate the unique features of each input data type we produced two primary assemblies. First, we assembled the short-read shotgun dataset using the heterozygosity aware *de Bruijn* graph-based algorithm Meraculous (Chapman et al. 2011). As we are interested in assembling a single haploid genome sequence, we collapsed the program’s resulting diplotigs into a single haploid assembly (*i.e.* “diploid mode 1”). Second, we assembled the processed ONT reads using wtdbg2 (Ruan and Li 2019) (Table 2). As expected given the substantially larger input read lengths, we obtained a much larger contig N50 using this program, than in our short-read based primary assembly (Table 2).

**Table 2.**
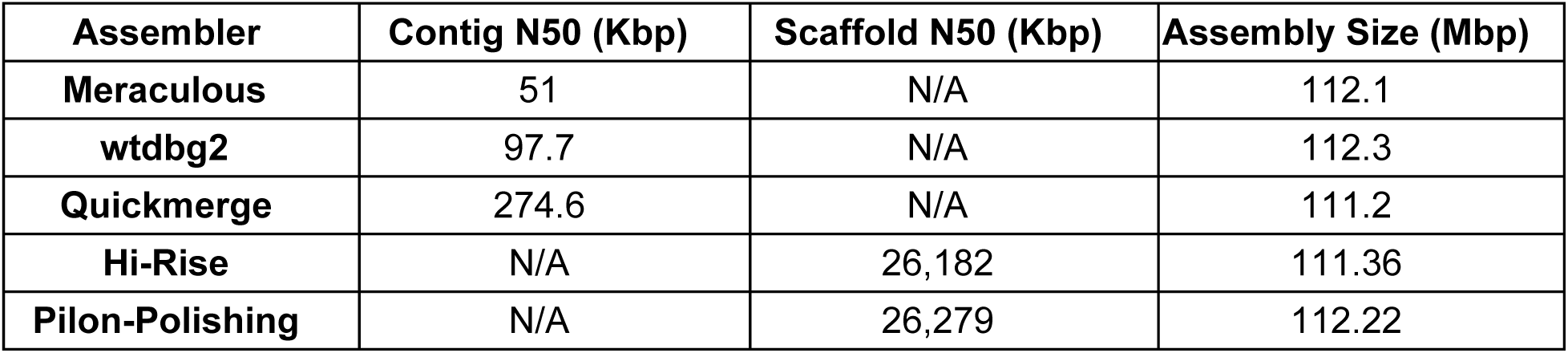
Summary of primary and scaffold assembly statistics.

### Merging Primary Assemblies

To combine the short and long-read primary assemblies we used the meta-assembler quickmerge. Quickmerge combines two input assemblies to produce an assembly with higher contiguity. Since the input assemblies come from the same individual, gaps in one assembly can be bridged by the other using the alignment of contigs from each input (Chakraborty et al. 2016). The resulting merged assembly had a contig N50 of 274.6 kb (Table 2).

### Scaffolding

Although the final merged primary assembly is reasonably contiguous, we observed by far the greatest gains in scaffold size after using our Hi-C data to scaffold the assembly (N50 = 26 Mb). Our final scaffolded assembly contains all the major chromosome arms in the *D. melanogaster* genome represented as single scaffolds, and correctly joins arms 2L and 2R across their heterochromatin-rich centromeric region (Figure 2). It therefore appears that the ability to produce high quality Hi-C libraries from extremely limited input material is the most essential component of our method for making contiguous genome assemblies for single individuals in small species.

**Figure 2:**
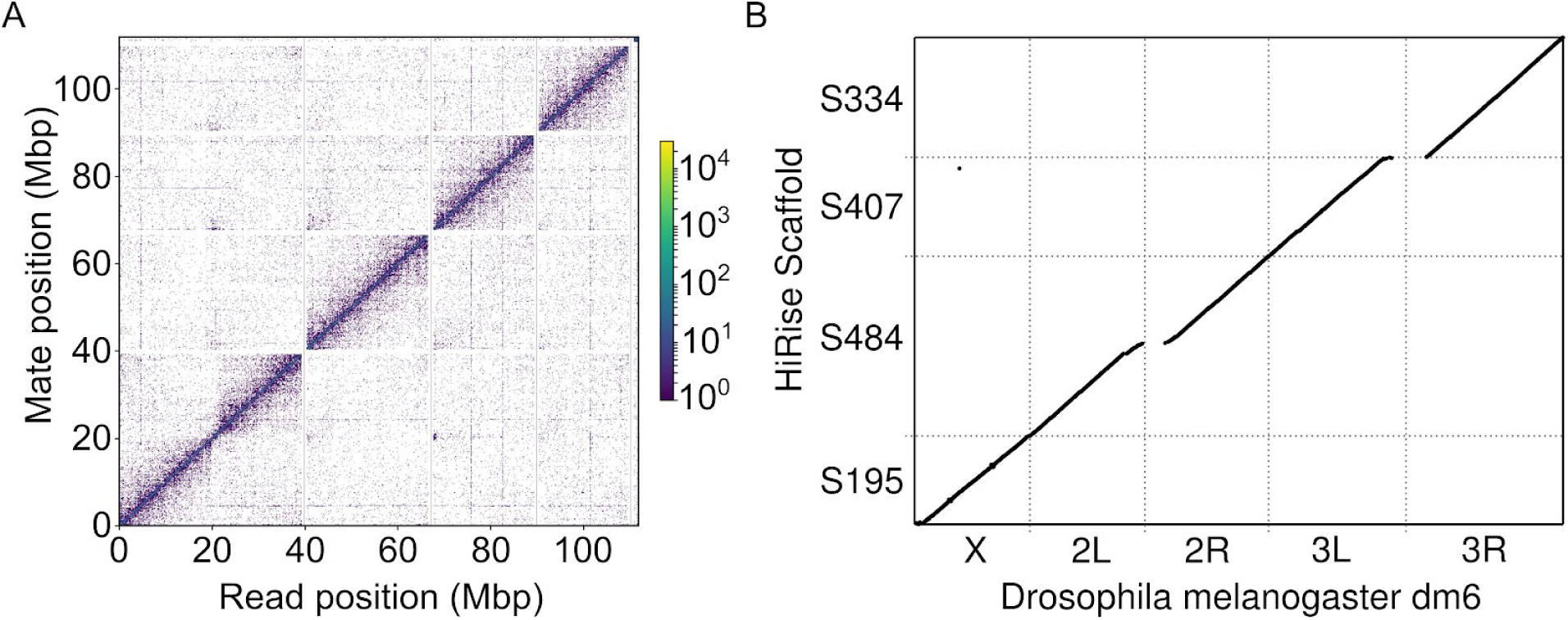
Genome Contiguity. (A) The read density map for Hi-C read pairs mapped onto the five largest contigs in our final assembly. (B) Dot plot of Hi-C scaffold assembly mapped to the dm6 reference genome. Continuous diagonal lines represent full length scaffolds of all major chromosome arms.

### Polishing and Gap Filling

Because we combined diverse data types, and in particular because our primary assembly relies on error-prone long reads, we sought to polish the contigs and fill gaps in the final highly contiguous assembly. In total we performed four rounds of iterative polishing with Pilon ((Walker et al. 2014), See Methods), until we did not observe significant additional improvements (Supplementary Table S2). The final assembly produced by this step, which we use for all validation below, is the largest of all of our assemblies at 112.22 Mb, which presumably reflects the success in our polishing and gap filling by incorporating additional sequences.

### Quality of the Final Assembly

We assessed our final assembly quality using several metrics. First, we applied the Benchmarking Universal Single-Copy Orthologs, BUSCO, algorithm (Simão et al. 2015). Briefly, the program provides an assessment of assembly quality specifically with respect to genic sequences by searching for a set of nearly-universal and single copy genes. In applying this quality metric we obtained a BUSCO score of 95.2% completeness for our final assembly. This is slightly lower than the current *D. melanogaster* ISO1 reference BUSCO score of 98.9%, but it is not dramatically different. We therefore conclude that the majority of the expected genic sequences are complete in our assembly.

Second, to compare the assembly of our H3 fly to the dm6 reference and quantify misassemblies we used the genome quality assessment tool QUAST (Gurevich et al. 2013). In addition, we used QUAST to compare another high quality assembly of a different *D. melanogaster* strain, A4 (Chakraborty et al. 2018), to the dm6 reference to set a benchmark for the expected differences between genetically diverse strains (Table 3). Because A4 was completely inbred and independently isolated from ISO1, whereas our H3 sample is heterozygous for the ISO1 genome, our assembly should more closely match the reference genome. The reason is that we would expect the reference allele to be selected 50% of the time at non-reference sites, and we should therefore observe approximately half as many apparent differences in our final assembly as for A4 relative to the ISO1 reference genome. As expected, our assembly had substantially fewer misassemblies, mismatches and indels than the A4 strain when compared to the dm6 reference, likely because of the relatedness between ISO1 and our assembled individual.

**Table 3.** Summary of QUAST output comparing H3 and A4 assemblies to the dm6 reference genome

|  | H3 against dm6 reference | A4 against dm6 reference |
| --- | --- | --- |
| # misassemblies | 798 | 2309 |
| # misassembled contigs | 15 | 145 |
| # local misassemblies | 1251 | 3491 |
| # mismatches per 100 kbp | 525.36 | 1136.97 |
| # indels per 100 kbp | 88.7 | 118.84 |

### Repeat Content

Despite similar BUSCO scores and the modest rate of misassemblies that we observe, our genome assembly is approximately 20% smaller than the canonical *D. melanogaster* reference genome. We suspected that much of the difference occurs because our assembly relies on relatively short reads and therefore collapsed repetitive regions. To evaluate this, we used the dm6 annotation data to evaluate coverage across different types of genomic features for both our single-fly assembly and a separate comparison of the A4 assembly. We found that while unique sequence including genes and especially exon sequences were captured in their entirety the majority of the time, highly duplicated elements such as transposons and tRNAs were much less likely to be covered by the H3 assembly (Table 4). This is a general weakness of short-read assemblies (Treangen and Salzberg 2011) and should be acknowledged by any forthcoming analysis applying this method of assembly.

**Table 4.** Sequence uniqueness strongly impacts assembly coverage. The columns are H3 assembly without any polishing and a non-reference control assembly of standard coverage and size. The rows are annotation types. The value corresponds to the percent of aligned annotated elements with at least 90% of their sequence captured in our assembly. The coverage distribution of our assembly is bimodal, with the vast majority of elements being either covered by a single assembled contig or not covered at all. An expanded table including more annotation types and counts, polished versions of the assembly, and overall assembly statistics can be found in the supplement (Supplementary Table S1).

|  | H3 Assembly | A4 Assembly Control |
| --- | --- | --- |
| **Coding Sequence (CDS)** | 94.0% | 97.9% |
| **Exon** | 93.9% | 99.5% |
| **Long noncoding RNA** | 90.6% | 98.8% |
| **microRNA** | 93.7% | 99.6% |
| **tRNA** | 76.5% | 98.7% |
| **Mobile genetic elements** | 55.3% | 82.0% |

### Phasing

We next evaluated our prospects for phasing the genome of this outbred individual, *i.e.* assigning each heterozygous allele to a chromosome. To do this, we realigned our short-read data to our final genome assembly and called all heterozygous variants using GATK (McKenna et al. 2010). We then realigned the Hi-C and long-read data as well and attempted to infer the phase using combinations of these data and the Hapcut2 algorithm (Edge et al. 2017). Because our individual is outbred and we know the complete genome sequence of both ancestors, it is straightforward to quantify the phase accuracy.

Using just the short-read data to phase heterozygous SNPs in the H3 individual, we achieve a modest combined mismatch and switch error rate (*sensu* (Edge et al. 2017)) of 0.00147 errors/site. Briefly, mismatch errors denotes sites where single variants are phased incorrectly in an otherwise correct block and switch errors denote a change where at least two subsequent variants are phased incorrectly relative to preceding sites. However the mean phase block length is just 14 heterozygous variants or approximately 2 kb. When we incorporated our Hi-C data, the combined error rate increased to 0.0147 error/site, but nearly entire chromosomes’ variants were included in a single phase block (*i.e*., 99.95% of variants/chromosome). The addition of Hi-C increased switch errors in particular by 0.0126 errors/site. This is likely a consequence of somatic chromosome pairing in dipterans (Cooper 1948), which has previously been demonstrated to create an excess of sister chromosome contacts in Hi-C data (Corbett-Detig et al. 2019; AlHaj Abed et al. 2019). The increased switch error rate suggests that approximately 17% of Illumina-phasable blocks that are joined by the addition of Hi-C result in switch errors. Therefore, phase inferred from these data could be useful across relatively short distances (*e.g.,* 5 kb), but should be regarded with caution at larger genomic distances. This might not be suitable for all applications of phasing, but would be sufficient for many population genetic questions that rely on short-distance haplotype and linkage information.

### Genomic Bycatch

Although not a primary consideration in this work, we found that our assembly captures additional material that is potentially of interest and underscores the power of our approach. First, our selected individual was phenotypically female, nonetheless, we discovered a non-trivial rate of Y-chromosome mapping contigs. Importantly, we found a similar Y-mapping rate in all three raw sequencing datasets (Supplementary Table S3), and the relevant Y:Autosome depth closely resembles that of typical phenotypic males (unpublished data). We therefore believe this is an XXY female. Despite the abundance of Y-derived reads, our Y chromosome assembly is exceedingly fragmented, as most Y chromosome assemblies are, reflecting the challenges of assembling extremely repeat-dense chromosomes (Kuderna et al. 2019). Nonetheless, this finding highlights the value of sequencing individuals rather than pools because pooling would likely obscure this relationship of relative chromosome depths.

Second, the reference strain is known to harbor the symbiotic bacteria *Wolbachia*, as we used this as the female parent in the cross *Wolbachia* is present in our sample due to infected embryos. Despite the differences in read-depths relative to the nuclear genome, our assembly includes nearly full coverage of the *Wolbachia* genome with few apparent misassemblies (Figure 3 and Supplementary Figure S2). *Wolbachia* in particular (Pietri et al. 2016), and endosymbionts more generally (Russell et al. 2019), are frequently present in host somatic tissues, likely explaining the similar abundances of *Wolbachia*-derived reads across sequencing libraries prepared from different parts of the fly. This suggests that in addition to nearly complete nuclear genomes, our assembly method might also be a powerful tool for investigating individual’s endosymbiont communities – a fundamental consideration in arthropod biology (Blow and Douglas 2019). Additionally, the analysis of a single individual obviates important concerns about pooling for interpreting inter-strain endosymbiont diversity (as in, *(Medina et al.*)), and again emphasizes the potential impact of this approach.

**Figure 3.**
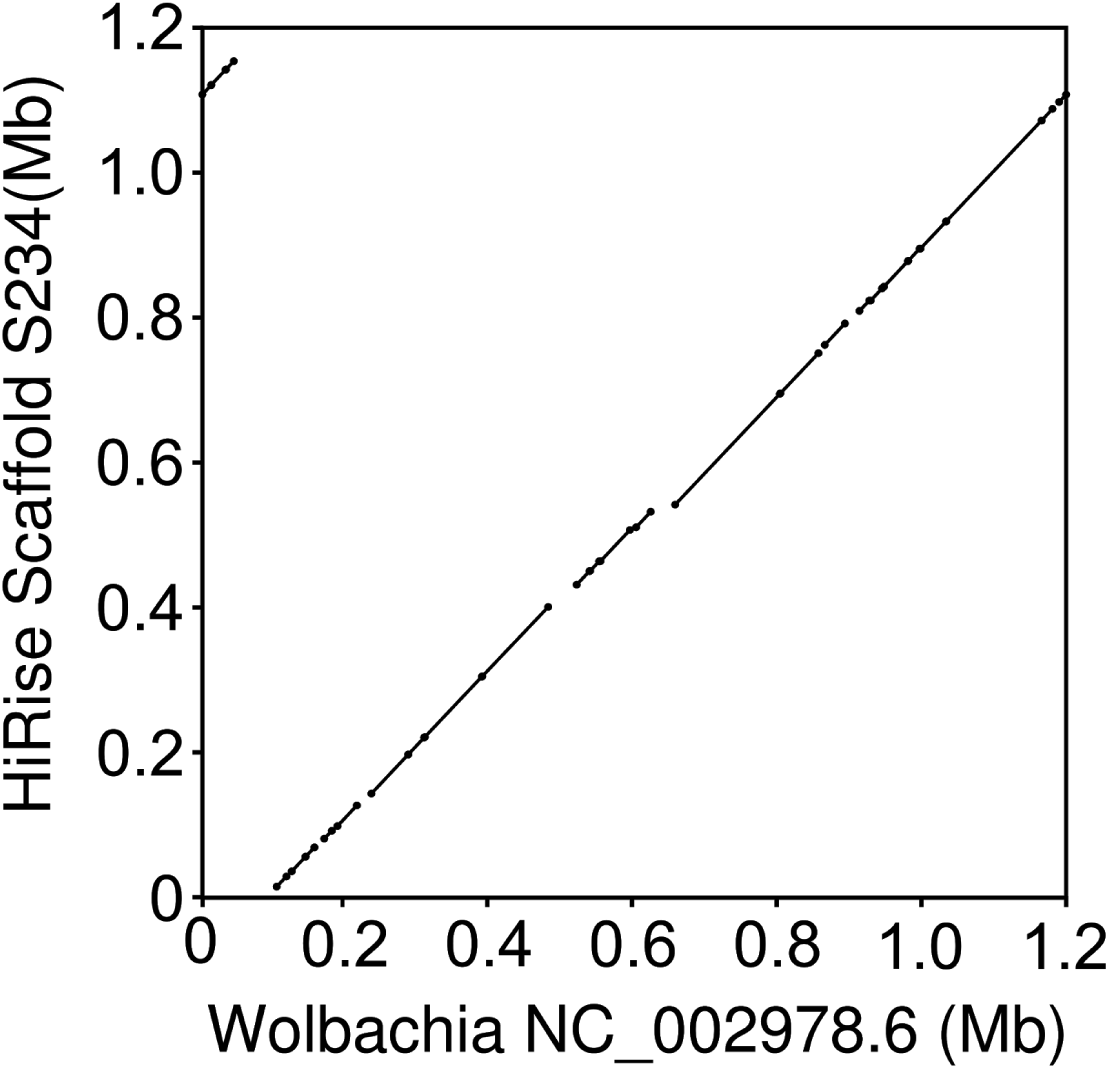
Dot-plot comparison of our nearly-complete *Wolbachia* assembly to the the canonical wMel *Wolbachia* genome sequence. Note that the apparent discontinuity in the top right/left, reflect the circular nature of the bacterial genome, and simply indicates that our assembly breaks the circle at a slightly different place.

## Conclusion

Recent advances in technology have greatly increased the quality of genome assemblies but generally require a relatively large DNA input. This limitation reduces the applicability of these methods for many precious, rare, and/or field collected specimens. Here, from a single fly we were able to construct a chromosome scale genome assembly with an N50 of 26 Mb. The primary assemblies were made with less than 90 ng of total input DNA. Therefore, our approach demonstrates that high quality chromosome-scale assemblies can be obtained from limited sample inputs.

Our method also compares favorably for total cost outlay. The DNA isolation and library preparation involves only basic molecular biology methods and equipment. We produced all necessary sequencing data on approximately one half of a HiSeq 4000 lane and a single MinION flow cell. We can therefore produce a contiguous, high quality genome for approximately $1,200 in total materials and reagent costs. For cost effectiveness, our approach compares quite favorably with available alternatives such as Pacbio SMRT cells at $2,000 each.

There are many genome assembly approaches available, and ours may not be optimal for all applications. When input materials are severely limited, the approach we describe here provides an appealing set of trade-offs and may be the only option to produce highly contiguous genome assemblies. Indeed, we have been able to make R2C2 libraries with as little as 10 ng of input DNA. Nonetheless, if more DNA is available, recent advances in PacBio library preparations (Kingan et al. 2019) might be a more appealing option for the long-read assembly. This method does not require amplification, and results in a less biased coverage. However, without Hi-C data for scaffolding, chromosome-scale assemblies are unlikely to be achievable. We therefore consider the addition of our Hi-C approach a necessary prerequisite for high quality genomes.

Perhaps the most fundamental concern for the suitability of our approach is the researcher’s specific questions and motivations for making a genome. Applications that require high contiguity in an assembly would be enhanced significantly using this approach. For example, association studies and quantitative trait locus mapping approaches generally require knowledge of large-scale linkage among sites to be successful (Ashton et al. 2017). Similarly, many population genetic frameworks, e.g. those for local ancestry inference (Maples et al. 2013; Corbett-Detig and Nielsen 2017), and for estimating past effective population sizes (Li and Durbin 2011), are based on the spatial distribution of markers along a reference genome. Finally, comparative studies of large-scale chromosome structure would be significantly enhanced by contiguous genome assemblies (Corbett-Detig et al. 2019). However, if the distributions of repetitive elements across the genome are of interest, our specific method is unlikely to perform well. Many studies are concerned primarily with coding regions, and for those our approach presents a reasonably high quality option.

This approach can serve as a guide point for genome projects of small organisms which make a large majority of the diversity of life. Approximately 80 percent of known species are insects, and approximately 5 million total insect species are believed to exist on earth (Stork 2018). Additionally, any research projects dealing with minimal DNA could achieve chromosome scale genomic information from this approach. This approach is therefore positioned to revolutionize our understanding of genome structure across diverse species.

## Materials and Methods

### DNA Extraction

High molecular weight DNA was extracted from one half of a single *Drosophila melanogaster* female using a Qiagen MagAttract HMW DNA kit. One half of a single fly was placed in a 1.5 ml tube with lysis buffer and proteinase k then crushed with a pestle using an up and down motion as to not shear DNA. The lysis and proteinase k digestion was incubated overnight at 37 C. The rest of the purification was performed according to the manufacturer’s protocol. The total amount of DNA recovered was 104.4 ng measured with a Thermo Fisher Qubit fluorometer and Qubit dsDNA HS assay kit. This sample was subsequently used for the Tn5 and nanopore library prep.

### Illumina Short-Insert Tn5 Sequencing

From the HMW DNA sample, 10 ng of gDNA was tagmented with Tn5 transposase for 8 minutes at 55°C. The reaction was halted by adding 0.2% SDS and incubated at room temperature for 7 minutes. Four separate PCR reactions were set up using the KAPA Biosystems HiFi Polymerase Kit and amplified for 16 cycles using uniquely indexed i5 and i7 primers. The amplified libraries were pooled and purified using the ≥ 300 bp cutoff on the ZYMO Select-a-Size DNA Clean and Concentrator Kit. 500 ng of the purified library pool was run on a Thermo Fisher 2% E-Gel EX Agarose Gel and cut between 550 and 800 bp. The gel cut was purified with the NEB Monarch DNA Gel Extraction Kit and quantified using the Qubit dsDNA HS Assay Kit and the Agilent TapeStation.

### Nanopore Sequencing

From the HMW DNA sample, 78.3 ng was used as input. The sample was first sheared using a Covaris g-TUBE centrifuged for 30 seconds at 8600 RCF. The sheared DNA was size selected using Solid Phase Reversible Immobilization (SPRI) beads at 0.7 beads:1 sample ratio and eluted in 25 ul ultrapure water.

End repair and A-tailing was performed using NEBNext Ultra II End Repair/dA-Tailing Module followed by ligation of Nextera adapters using NEB Blunt/TA Ligase Master Mix following the manufacturer’s protocol. The adaptor ligated sample was purified by SPRI beads at a 1:1 ratio and eluted in 50 ul of ultrapure water. The sample was divided into six, 25 ul PCR reactions with Nextera primers and KAPA HiFi Readymix 2x (95 C for 30 s, followed by 12 cycles of 98 C for 10 s, 63 C for 30 s 72 C for 6 min, with a final extension at 72 C for 8 min then hold at 4 C). The PCR reactions were pooled and purified by SPRI beads at a 1:1 ratio and eluted in 60 ul of ultrapure water. Concentration was measured to be 110 ng/ul using the Qubit dsDNA HS assay. The entire sample was size selected by gel electrophoresis using a 1% low melting agarose gel. An area from 6-10 kb was cut out and digested using NEB Beta Agarase I following the manufacturer’s protocol then purified using SPRI beads at a 1:1 ratio.

One hundred nanograms of size selected DNA was mixed with 50 ng of a DNA splint and circularized by Gibson assembly using 2x NEBuilder HiFi DNA Assembly Master Mix incubated for 60 min at 50 C. Non circularized DNA was digested overnight at 37 C using Exonuclease I, Exonuclease III and Lambda Exonuclease (all NEB). Circularized DNA was purified by SPRI beads at a 0.8:1 ratio and eluted in 40 ul of ultrapure water.

The circularized DNA was split into 8 50 ul rolling circle amplification (RCA) reactions (5 ul 10x Phi29 buffer (NEB), 2.5 ul 10 mM dNTPs (NEB), 2.5 ul 10 uM exonuclease resistant random hexamer primers (Thermo), 5 ul DNA, 1 ul Phi29 polymerase (NEB), 34 ul ultrapure water). Reactions were incubated overnight at 30 C. All reactions were pooled and debranched using T7 Endonuclease (NEB) for 2 hours at 37 C. To shear ultra-long RCA products the sample was run through a Zymo Research DNA Clean and Concentrator-5 column and eluted in 40 ul ultrapure water. A final size selection was performed by gel electrophoresis using a 1% low melting agarose gel. An area at approximately 10 kb was cut out and digested using NEB Beta Agarase I following the manufacturer’s protocol then purified using SPRI beads at a 1:1 ratio.

The cleaned and size selected RCA product was sequenced using the ONT 1D Genomic DNA by Ligation sample prep kit (SQK-LSK109) and a single MinION flow cell following the manufacturer’s protocol. The raw data was basecalled using the Guppy basecaller. Consensus reads were generated by Concatemeric Consensus Caller with Partial Order alignments (C3POa).

### HiC Library

The anterior half of the fly was placed into a 1.5 ml tube with 1 ml of cold 1x PBS. 31.25 ul of 32% paraformaldehyde was added. The sample was briefly vortexed and incubated for 30 minutes at room temperature with rotation. After incubation the supernatant was removed and washed twice with 1 ml of cold 1x PBS. 50 ul of lysate wash buffer was added before grinding with pestle. 5 ul of 20% SDS was added then vortexed for 30 seconds and incubated at 37 C for 15 minutes with shaking. 100 ul of SPRI beads were added to bind chromatin. Bound sample was washed 3 times with SPRI wash buffer.

Beads were resuspended in 50 ul of Dpn II digestion mix (42.5 ul water, 5 ul 10x DpnII buffer, 0.5 ul 100 mM DTT, 2 ul DpnII) and digested for 1 hour at 37 C with shaking. Beads were washed twice with SPRI wash buffer and resuspended in 50 ul of end fill-in mix (37 ul water, 5 ul 10X NEB Buffer 2, 4 ul 1 mM biotin-dCTP, 1.5 ul 10 mM dATP dTTP dGTP, 0.5 ul 100 mM DTT, 2 ul Klenow fragment) then incubated for 30 minutes at room temperature while shaking. Beads were washed twice with SPRI wash buffer and resuspended in 200 ul of intra-aggragete mix (171 ul water, 1 ul 100 mM ATP, 20 ul 10x NEB T4 DNA Ligase Buffer, 1 ul 20 mg/ml BSA, 5 ul 10% Triton X-100, 2 ul T4 DNA ligase) then incubated at 16 C overnight while shaking. Beads were placed on a magnet to remove supernatant then resuspended in 50 ul of crosslink reversal buffer (48.5 ul crosslink reversal mix, 1.5 ul proteinase K) then incubated for 15 minutes at 55 C, followed by 45 minutes at 68 C while shaking. Beads were placed on a magnet and the supernatant was transferred to a clean 1.5 ml tube. 100 ul of SPRI beads were added to the supernatant and allowed to bind before washing twice with 80% ethanol and eluting sample with 50 ul of 1X TE buffer.

The sample was then fragmented by sonication. Fragmented sample was end repaired and adapter ligated using the NEBNext Ultra II kit following the manufacturer’s protocol. The sample was purified from ligation reaction by SPRI beads, washed twice with 80% ethanol, and eluted in 30 ul of 1X TE. Biotin tagged fragments were enriched using streptavidin C1 Dynabeads. Enriched fragments were indexed by PCR (23 ul water, 25 ul 2x Kapa mix, 1 ul 10 uM i7 index primer, 1 ul 10 uM i5 index primer) and amplified for 11 cycles. Reaction was purified by SPRI beads and quantified using the Qubit dsDNA HS Assay Kit and the Agilent TapeStation.

### Assembly

We produced short-read assemblies using the variation-aware *de Bruijn* graph algorithm, Meraculous (Chapman et al. 2011). Long-read data was assembled using Wtdbg2 (Ruan and Li 2019) using the following options “wtdbg2 -x ont -g 120m -p 0 -k 15 -S 1 -l 512 -L 1024 --edge-min 2 --rescue-low-cov-edges” followed by the wtdbg2 consensus caller wtpoa-cns (Ruan and Li 2019). The two primary long and short-read assemblies were combined using quickmerge default merge_wrapper.py command.

### Scaffolding

We polished the hybrid shotgun and long-read assembly using the Illumina shotgun dataset using the bwa mem algorithm (version 0.7.17) (Li and Durbin 2009) to map the Illumina reads back to the genome and samtools (version 1.7) to sort the reads. We input the sorted alignment to the consensus for wtdbg (wtpoa-cns) (version 2.5) using the command “-x sam-sr” to polish the contigs of the hybrid assembly. We scaffolded the polished assembly using the scaffolding tool HiRise (version 2.1.1) run in Hi-C mode using the default parameters with the Hi-C library as input.

### Polishing

The draft assembly went through a total of four iterative rounds of polishing using the automated software tool Pilon using default settings. For each round the short and long-read data was mapped to the draft assembly using minimap2. After each round, the assembly was evaluated for misassemblies, indels, mismatches, N50, and assembly size using QUAST (Gurevich et al. 2013) to determine if further polishing would increase the assembly correctness.

### Evaluation

To evaluate the completeness of the H3 assembly we searched for conserved genes using Benchmarking Universal Single-Copy Orthologs v3, (BUSCO) with the metazoa odb9 lineage gene set (Simão et al. 2015). To compare to the current reference genome we used the genome quality assessment tool QUAST using the “--large --k-mer-stats” options (Gurevich et al. 2013). Misassemblies are defined by the following criteria, a position in the assembled contigs where 1) the left flanking sequence aligns over 1 kbp away from the right flanking sequence on the reference, 2) flanking sequences overlap on more than 1 kbp, 3) flanking sequences align to different strands or different chromosomes. Local misassemblies are defined by the following criteria 1) the gap or overlap between left and right flanking sequences is less than 1 kbp, and larger than the maximum indel length (85 bp), 2) The left and right flanking sequences both are on the same strand of the same chromosome of the reference genome.

### Repetitive and genic region coverage analysis

We aligned three separate versions of H3 assembly with zero, one, and two rounds of polishing with Pilon to the *Drosophila melanogaster* reference using Minimap2 with default parameters and sam output (Li 2018; Walker et al. 2014). We then applied samtools compression and sorting to produce sorted bam files ((Li et al. 2009; Quinlan and Hall 2010)), to which we applied bedtools genomecov with options -ibam and -bga to produce a file of region coordinates and coverage values of 0 or more for each region across the genome (Li et al. 2009; Quinlan and Hall 2010). We combined this information with the annotation gff3 file with a custom script that assigned coverage values to all annotated spans base by base (Quinlan and Hall 2010). The average coverage per base was calculated for each annotated span, then the average and mean value of coverages for all spans for each annotation type was calculated. As a control for comparison we performed this procedure on a complete non-reference *melanogaster* assembly and calculated similar values to elucidate any particular weakness our assembly exhibits.

### Phasing

To phase the genome, we realigned all short-read data to our final genome assembly using BWA mem (Li and Durbin 2010). We then called all heterozygous variants using GATK (McKenna et al. 2010) on the four largest scaffolds in our assembly, and we filtered this set to exclude SNPs and indels in the bottom 10% or top 10% of observed sequencing depths. As the H3 genome is a mosaic of I38 and dm6 alleles, we “polarized” each heterozygous variant by realigning the dm6 genome using minimap2 (Li 2018) to determine whether H3 contained the dm6 allele. We then aligned all Hi-C data using BWA mem (Li and Durbin 2010) and the ONT data using minimap2 (Li 2018) and attempted to phase the genome using varying combinations of these data using hapcut2 (Edge et al. 2017). We quantified mismatch and switch errors as described in (Edge et al. 2017).

## Data Access

The sequencing data generated in this study has been submitted to the NCBI BioProject database (https://www.ncbi.nlm.nih.gov/bioproject/) under accession number PRJNA591165.

## Acknowledgments

This work was supported by the NIH (R35 GM128932 to RBC-D and R35 GM133569-01 to CV) and from an Alfred P. Sloan Fellowship to RBC-D. During this work JM was supported by NIH training grant T32 HG008345-01.

